# Wood ants express uncertainty when engaging with magnetic cues for directional guidance

**DOI:** 10.1101/2021.04.18.440333

**Authors:** Thomas S. Collett, Andrew O. Philippides

## Abstract

Wood ants were trained indoors to follow a route in a chosen magnetic direction from the centre of a small, circular arena to find a drop of sucrose at the edge. The arena, surrounded by a white cylindrical wall, was in the centre of a 3D coil system that generated an inclined Earth strength magnetic field in any horizontal direction. Between trials, the chosen magnetic training direction was rotated to a new orientation. Tests were given without food and with fresh or reversed paper on the floor of the arena. In a significant number of tests, ants left the centre facing the goal, or in the opposite direction, but they mostly failed to reach the goal. Tests given early in the day, before any training, show that ants remember the magnetic route direction overnight. On some training trials, the position of the sucrose was also indicated by a black stripe. Not uncommonly, ants first moved in the opposite direction to the stripe before switching to the correct direction. Travel away from the reward seems to express the ant’s uncertainty about the correct path to take. Tests show that this uncertainty may stem from competing directional cues linked to the room, suggesting that ants are reluctant to rely on magnetic information alone. We conclude that ants can remember a route direction defined by an Earth-strength magnetic field and that they express any uncertainty about the correct direction by moving for a stretch in the opposite direction. In a second experiment, an upright and an inverted triangle were fixed 90° from each other to the inside of the cylinder. Sucrose was placed beneath one of the triangles, dependent on the direction of the magnetic field. Ants failed to master this task and to approach the magnetically cued triangle. Instead, they preferred to approach the upright triangle. The ants were again uncertain of the correct direction and expressed this uncertainty through paths that had segments directed towards both the inverted and the upright triangles.

## INTRODUCTION

Ants and bees can guide their paths during navigation using magnetic cues (reviews: Wajnberg et al., 2010, Wiltschko, 2012, Fleischmann et al. 2020). An excellent example comes from the desert ant, *Cataglyphis noda* (Fleischmann et al. 2018): young *C. noda*, before their sun compass is calibrated, rely on magnetic cues to provide compass information during their learning walks when they first leave their nest and are naïve to the world outside it (review: Zeil and Fleischmann, 2019). During the walks, ants loop around the nest, periodically turning towards it, giving themselves the opportunity to memorise views that can guide their later returns to the nest. Manipulation of the direction of the magnetic field during normal outdoor learning walks reveals that turns to face the nest are directed by path integration with the Earth’s magnetic field providing compass information. Desert ants do not seem to rely on magnetic information in later life and switch to a time compensated sun compass that may give more precise directional information (e.g. Wehner and Müller 2006). The low signal to noise ratio of magnetic information seems to be a general feature among animals (Johnsen et al 2020). Magnetic information may often be reinforced or rendered uncertain by other cues to direction (Johnsen et al 2020), as also happens in our ants.

The wood ant, *Formica rufa*, is known to be sensitive to the Earth’s magnetic field (Çamlitepe and Stradling, 1995). We examined whether these ants can remember the magnetic direction of a route to a food site. Ants were trained in a 3-D 1 m^3^ magnetic coil system to go in a specified magnetic direction from the centre of a small circular arena (38 cm diameter) to reach food at a point on the circumference. To prevent other potential directional cues from being persistently reinforced, the chosen magnetic direction was rotated after each trial by ca. 100° with the direction of rotation switched each day. In some trials, especially at the start of training, the food site was also indicated by a vertical black stripe placed on the arena wall just above the food. We report the ants’ behaviour during training trials with a stripe and during tests in the absence of sucrose and the stripe. These experiments emphasise that while ants are successful in learning the magnetic direction of a route, they are nonetheless uncertain about following the magnetically signalled direction and express their uncertainty by moving in the opposite direction to that signalled by the magnetic cue (See Discussion).

A second series of experiments explored whether magnetic cues to direction can provide a contextual cue that enables ants to select between two routes. Ants were released at the centre of the same arena and were confronted with an upright and an inverted triangle in fixed positions on the arena wall. Sucrose was placed at the bottom of one of the triangles depending on the orientation of the magnetic field in the arena. Previous work suggests that ants might be able to solve this problem. They distinguish between these shapes and can be trained to approach either of them (Judd and Collett. 1998). Indeed, honeybees can be trained to use magnetic direction to decide which of two patterns they should choose (Frier et al, 1996). In contrast to honeybees, the ants seemed unable to master this task, despite their ability to learn a magnetic direction.

## METHODS

### Ants

Experiments were performed on three colonies of laboratory maintained wood ants, two in 2018 and one in 2019. All the colonies were collected from Broadstone Warren, East Sussex, UK. Initial experiments from, January to March 2019, were on 2018 colonies that had been in captivity for at least 6 months. In the event, these initial experiments turned out to be a training experience for the experimenter, rather than for the ant, but the sparse data available from the experiments indicated that ants were responding to the magnetically specified direction.

Experiments on the 2019 colony took place between June and August, a few weeks after the colony was taken to the laboratory. All the presented data come from this colony. The colonies were kept under a 12 h:12 h light-dark cycle and were sprayed with water daily. Normally, water and sucrose dispensers were always available. During experiments, the colony was given limited access to sucrose to encourage enthusiastic foraging. Crickets were supplied several times a week.

Before training about 30 ants were marked individually with coloured enamel paint (Testor) and about a third of this group completed training.

### Magnetically controlled arena

Three pairs of 1 m diameter coils arranged in a cube, made by claricent, generated a uniform Earth strength inclined magnetic field within a volume of about 50 cm^3^. Three computer controlled power supplies (Tenma Model 72-2685 Digital Controlled DC Power Supply) and power amplifiers (constructed in house) determined the magnetic direction within the central cube in 5° steps. On each trial, the set directions were confirmed with a magnetic compass.

### Route learning

#### Ant training

Ants were trained to find sucrose in a specified cardinal magnetic direction on a paper covered horizontal board in the centre of the coil. A white painted cylinder (38 cm diameter) with its bottom removed was placed in the centre of the platform. Ants taken from their nest were released in the centre of the circular arena at the bottom of the cylinder and could find a drop of sucrose at the edge of the arena in the specified magnetic direction from the arena centre. After every trial, the magnetic direction was rotated by about 100° relative to the laboratory. Each day, the direction of rotation switched between clockwise and anticlockwise. To minimise the use of trail cues, the paper on which the ants walked was frequently shifted relative to the bucket and reversed or replaced.

The specified cardinal direction was changed between experiments. The 2018 colony was trained with magnetic directions, North, South and East. The presented data come from the 2019 colony trained with a route to the West.

Training was in two stages. In the first stage, individually marked ants were trained in groups of 5 to 6 individuals. A group of ants was put in a small release cylinder (7 cm wide, 5 cm high) with a 30° exit in the wall that pointed in the direction of the sucrose. At the start of training and periodically through training, the magnetic direction to the sucrose was reinforced by a black vertical stripe (6cm high by 0.5 cm wide) held by ‘Blu-Tack’ to the inner wall of the cylinder just above the sucrose.

In the second stage of training, ants were released individually in the centre of the arena from a portable release compartment. The device consisted of two parts: first, a Perspex cylinder (2.7 cm high, 2 cm wide) with an open bottom and closed top; second a Perspex circular base plate (5 cm wide) with a circular groove in its centre to accept the open end of the cylinder. A length of nylon fishing line was attached to the top of the cylinder. An ant was placed inside the cylinder with the open end facing upwards and the base plate put on top with the cylinder slotted into the groove. The release compartment was then placed right side up in the centre of the arena in no particular orientation. The fishing line was looped over a support 58 cm above the arena so that the cylinder could be pulled up vertically to a level just above the recording video camera and then secured - thereby releasing the ant. This method frees the ant to move in any horizontal direction. On a few occasions, a trial was aborted because the ant clung to the inside of the cylinder and was raised with it.

As training progressed, the stripe was removed on many of the trials. If an ant failed to reach the sucrose when there was no stripe, the stripe was replaced to ensure that the ant was rewarded. Because the arena is so small, it is difficult to exclude the possibility that, during training, ants are attracted directly to the drop of sucrose by cues emanating from it. Consequently, we only examined the ants’ behaviour during training, when that possibility did not affect the question that was asked.

An HD webcam (Microsoft 6CH-00002,1200 x 1080 pixels), which surveyed the whole arena from a position just above the coils recorded the ant’s path at 30 fps from the moment the cylinder was raised above the base plate. Paths were stored on computer as MP4 files that were later re-coded as AVIs and then processed with custom written code in MatLab to extract both position and facing direction of the ant. These data were checked and corrected by hand using methods described in de Ibarra et al., 2009.

#### Tests

Once ants were trained, 9 trials were given with the stripe and sucrose missing. Before each test, the bucket and magnetic field were rotated in opposite directions and the paper surface under the bucket was changed or turned over. To see whether ants have a ‘longer’ term memory of the magnetic direction, 4 tests were given at the start of the experimental day before there had been any training. During training and tests not all ants in the cohort could be found on the surface of the nest. We include all the test trajectories of 9 ants that appeared for at least 6 of the 9 tests that were given.

#### Data presentation

Because the ants’ paths were varied, the raw paths are shown. For tests, we show the initial parts of all the trajectories recorded from 2 of the 9 ants. In a significant number of cases, the ant, at the start of the test, faced the goal or a fictive goal in the opposite direction. These facing directions are illustrated in plots of facing direction against time from the start of a test.

A second measure is the overall direction of the ants’ path. The ant may walk in what looks like the correct direction for some seconds but then lose track and search. We decided subjectively when this change happens and cut the test path at this point. Path direction is assessed by plotting the distance that the ant travels in each direction, accumulated within 10° bins. This measure minimises the effect of periods where the ant is either still or moves very little when the direction of travel is unreliable. Since ants tend to travel in both directions towards a real and a diametrically opposite fictive goal, direction is given axially with 0° indicating movement in either direction along the magnetically defined axis.

Smoothing of the path is needed to remove distortions to the overall travel direction caused by the typical zigzag path of an ant to its goal (Lent et al., 2013). Thus, to calculate the ant’s direction of movement at each step, we first remove any points where the ant does not move (as angular direction is undefined at these points). We then smooth the x and y positions separately using a running average over 33 points (1 second), 16 before and 16 after the current data point (for the first/last 16 points, the data is padded by replicating the first/last point).

### Magnetic direction as a contextual cue

Ants were released in the centre of the arena as already described. They chose between approaching an upright or an inverted triangle on the cylindrical arena wall to find sucrose at the base of one, depending on the orientation of the magnetic field. The sucrose was beneath the upright triangle, when the direction from the central start point to the upright triangle was West. The inverted triangle was rewarding when the magnetic direction to that triangle was North. The triangles (7.8 cm high, 6.3 cm base) were cut from black card. They were fixed in position, 90° apart with their tip or base on the arena floor. The ants’ paths were recorded during training and test trials following the same procedure as in the route learning experiment. Ants were equally wrong during training and tests.

## RESULTS

### 1. Route Learning

#### (i) Training trials with visual cues

During training trials in which the ant is guided to the sucrose by the black stripe and by magnetic direction, the ant usually reaches the food (e.g., Figure 1). Unexpectedly, their approach to the sucrose was often preceded by a phase in which the ants move in the opposite direction. This effect is illustrated by the seven training trials from one ant in which the stripe was present. The ant travelled directly towards the stripe and the food in only one trial (trial 9). In three trials (11, 40, 42) the ant left the start and moved for a short stretch in the opposite direction. In three trials (34, 40, 42), the ant went all or most of the way to a fictive goal in the diametrically opposite position to the real goal. In two trials (19 and 44) the ant’s intentions in the first phase were not clear. Similar examples of travel in the opposite direction occur in other ants (Supplementary Figure 1). This behaviour cannot be attributed to some quirk in the applied magnetic field as it also occurred in three trials at the start of the day when the coils were inactive and the ants were guided by the Earth’s natural magnetic field (Trial 11 in Figure 1; 2 further cases in Figure S1).

**Figure 1.**
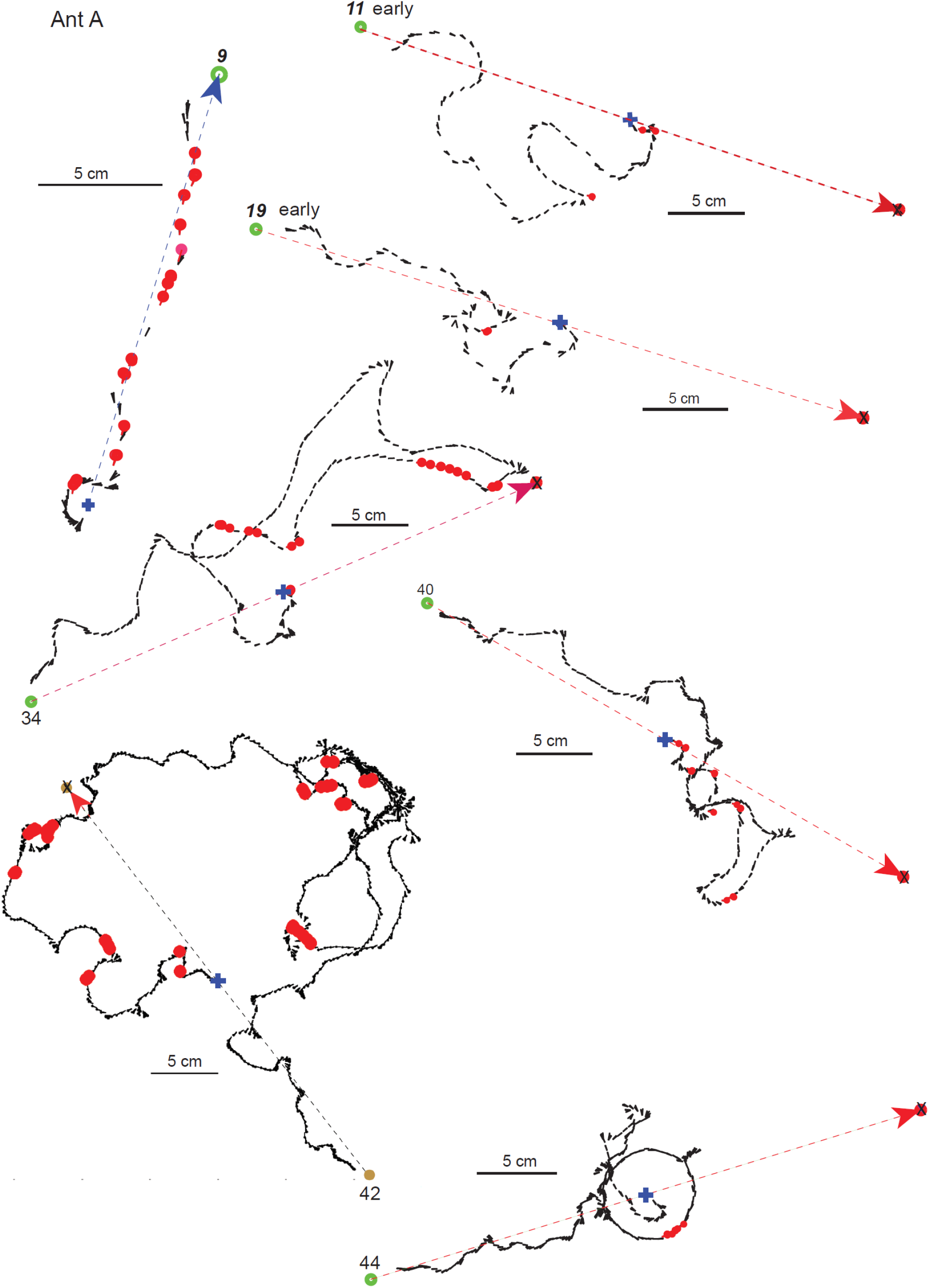
Training trials with magnetic direction supported by a black bar. Each panel shows the path of the same ant on different training trials with the position and orientation of the ant (black ball and stick) shown at 30 fps. Blue cross marks starting point. Red disks in the trace of the ant’s path indicate frames in which the ant faces (within +/− 10°) either towards the food (marked with a coloured disk) or towards a fictive goal in the opposite direction (coloured disk with an X). The real and fictive goals are both 18 cm from the central starting point. The colour and direction of the arrow between the start and goal(s) indicates whether the red disks show the ant facing the real (blue) or fictive (red) goal. Numbers by the goals indicate which trial the path is from. In trials marked ‘early’, i.e. early in the day, the coil system was off and the ant was guided by the Earth’s magnetic field.

**Figure 2.**
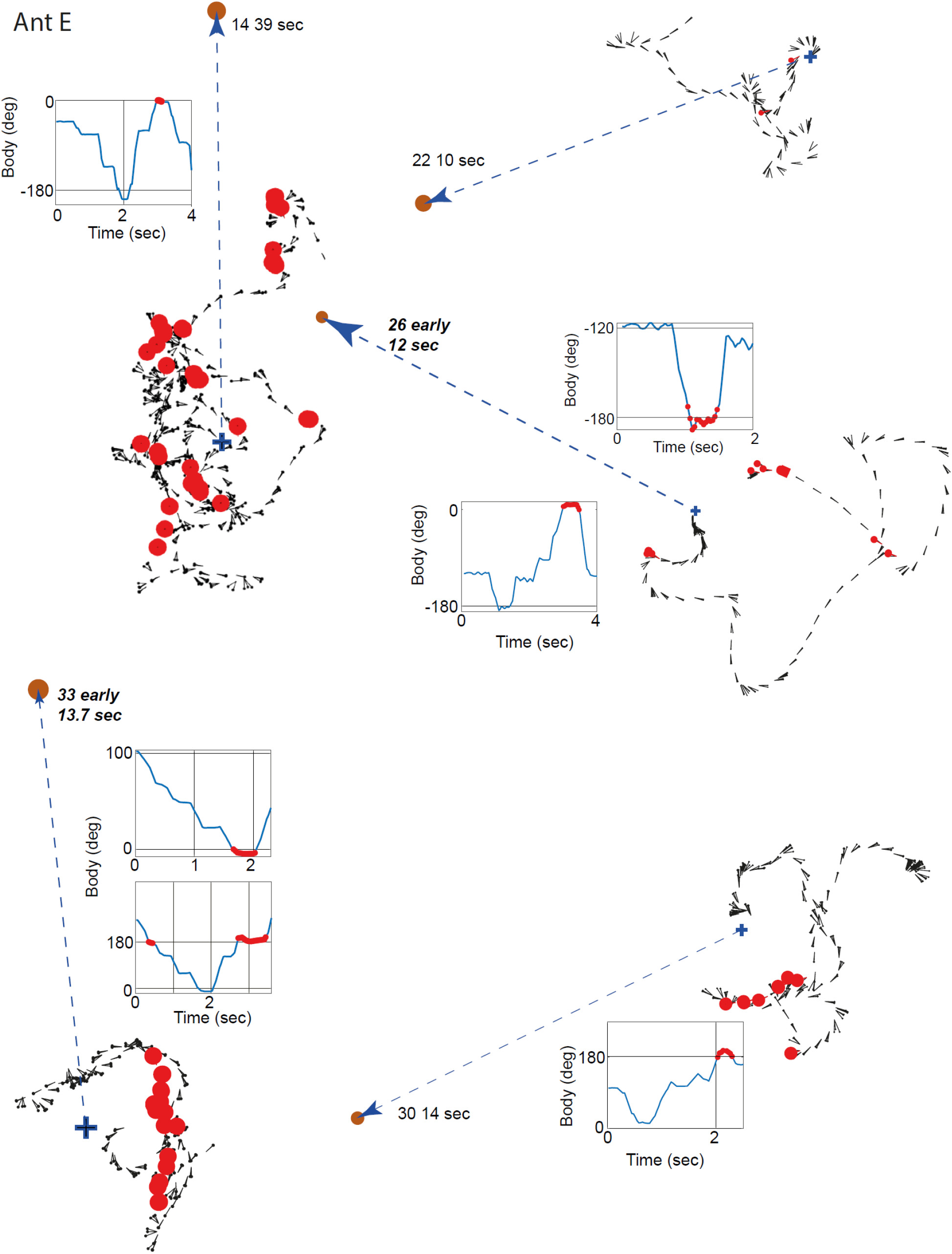
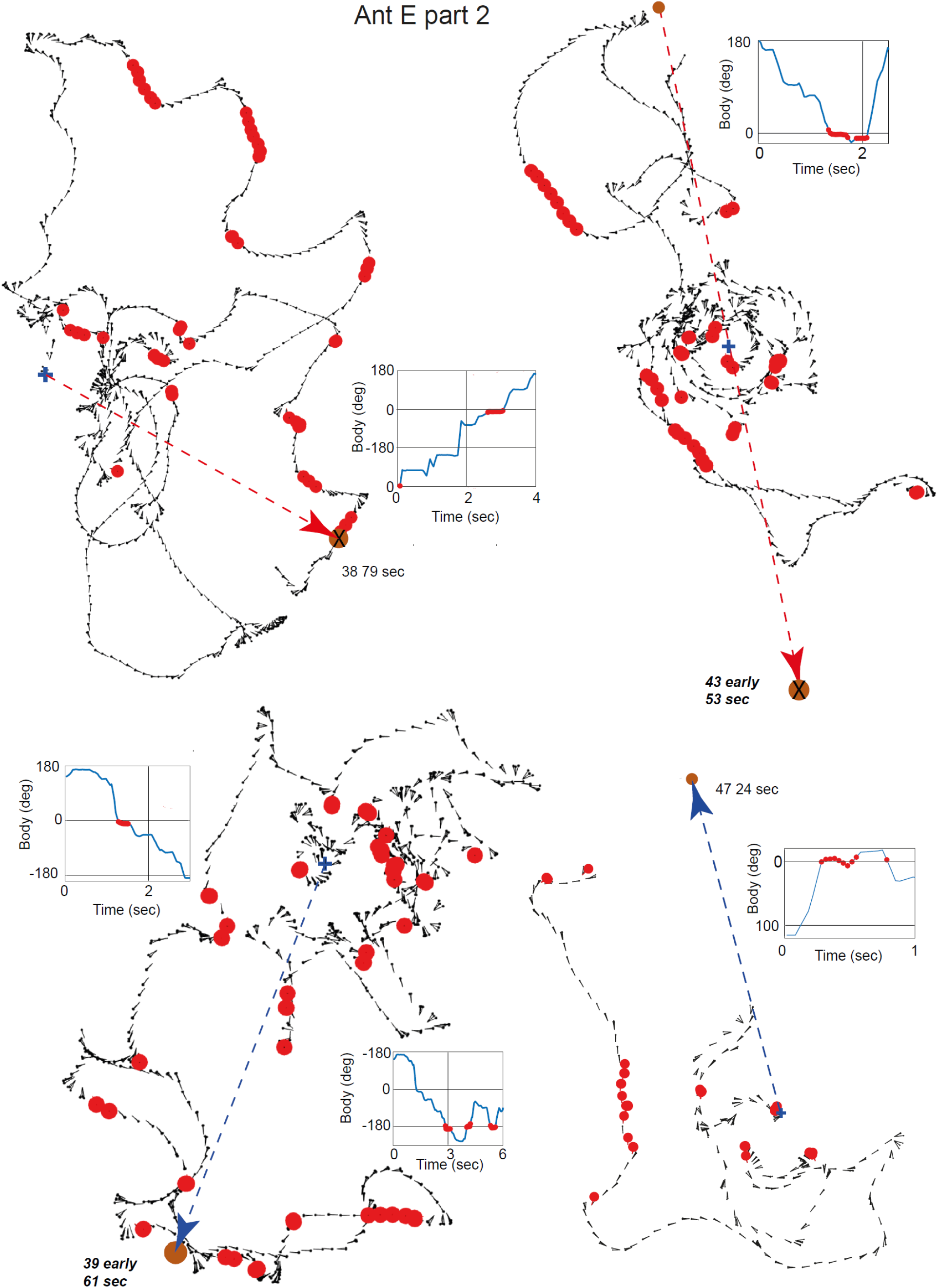
All the trajectories of ant E during its tests. Each panel shows part or all of a path recorded during one test, with the duration of the excerpt given below the test number. The orientation of the ant’s body over time relative to the magnetically defined goal is shown with 0° or 180” indicating that the ant faced the real (0°) or fictive (180°) goal. Red dots indicate that body orientation lies within ±10° of a goal. 0 sec is at the start of the path. ‘early’ indicates that the test was given before any training that day. Other details as in Figure 1.

**Figure 3.**
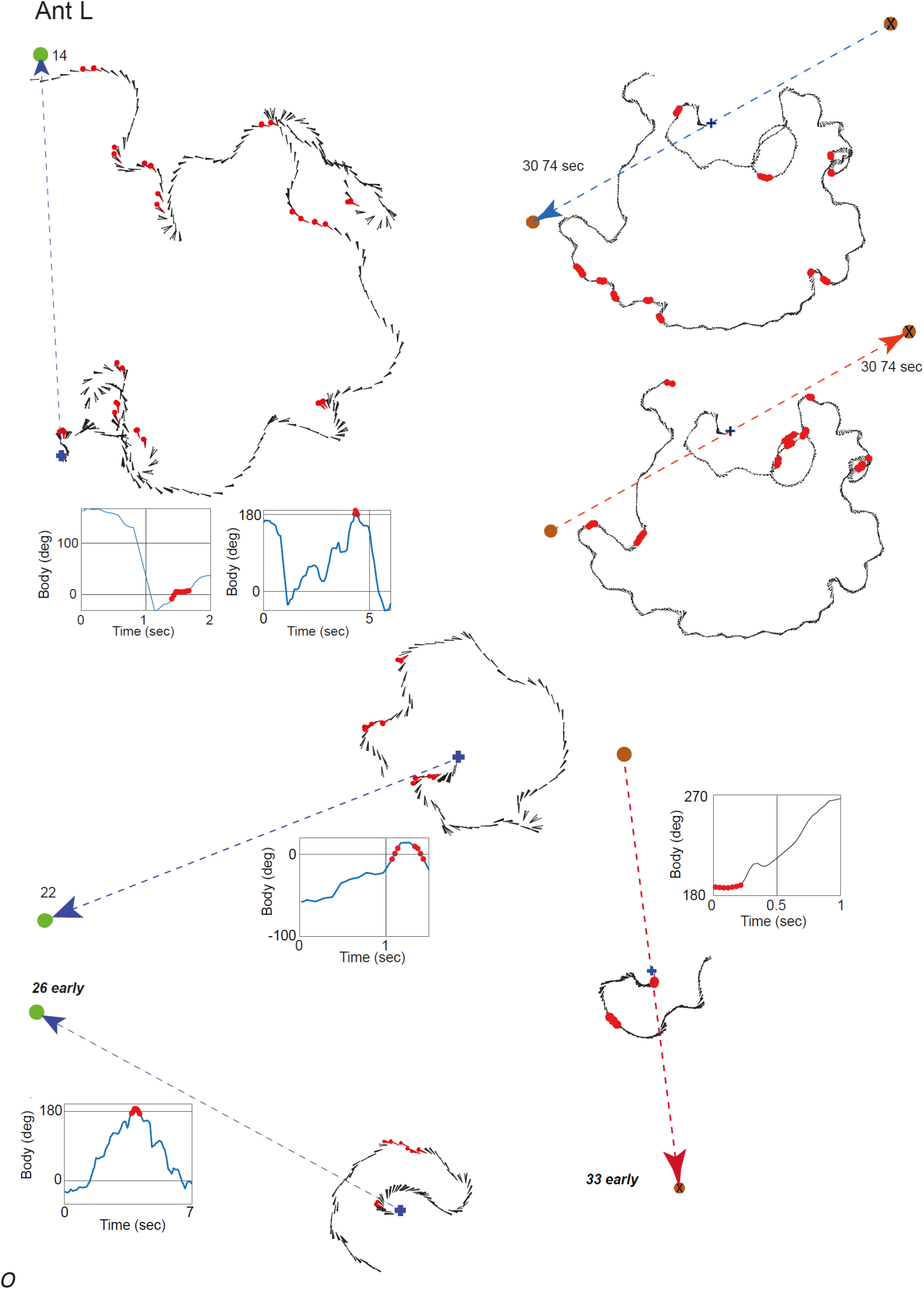
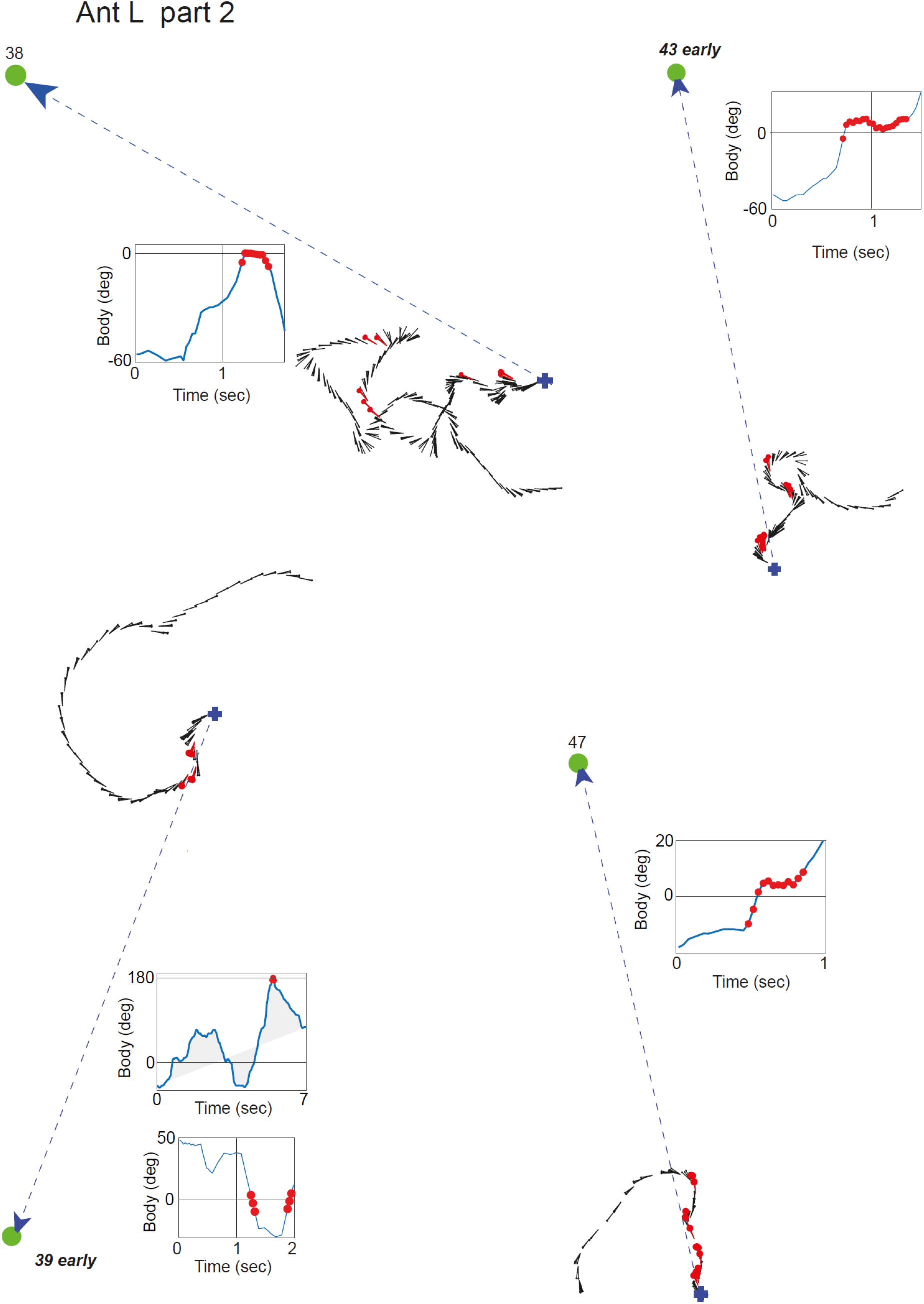
All the trajectories of ant L during its tests. Details as in Figures 1 and 2. Tests 22 and 30 were failures the other 6 were successful.

To assess the overall frequency of trajectories that started or were predominantly in the opposite direction to the stripe, we scored the 9 ants that contributed to 6 or more of the 9 tests, on the assumption that these were the most experienced ants. Out of a total of 52 training trials with the stripe, this sample of ants moved in the opposite direction to the stripe on 22 occasions, using the same subjective, but conservative, criteria as for the paths in Figure 1. The exact binomial test for a success of 22 or more trials on the conservative assumption their direction lay within ±40° (i.e. 80/360) of the opposite direction was significant (P =0.000447)

#### (ii) Tests

To give an idea of the diversity of ant tracks during tests, we show all the recorded tests of two ants. A significant proportion of trajectories have a stretch within the first 10 seconds of the test in which their body is oriented towards the real goal or the fictive goal lying in the opposite direction. Of the 4 early tests given before training, only in test 26 could ants the Earth’s natural field. In the other 3 tests, the direction of the goal was set by the applied magnetic field. To determine the number of tests in which ants faced in the correct or opposite direction at the start, we plotted the ant’s body orientation over time. A success was scored either when there is a substantial stretch close to the start in which the ant faced the real or fictive goal (±10°) as in tests 33, 43 and 47 of ant L, or when there was a short consecutive sequence of at least two 33ms video frames at a trough or a peak in the orientation-time plot in which the ant faced the real goal or fictive goal (±10°). The reason for requiring a trough or peak is that in these cases ants often face towards one of the goals and then switch and turn to face the other goal, as in tests 26 and 33 of ant E. It seems that when ants come to face in a goal direction, they have often reached a point of decision about where to face next.

We step through the tests of ant E to make the scoring clear. The following 6 tests are positive: 14, 26, 30, 33, 43, 47. The plots of tests 38 and 39 are taken as failures because, although there are plateaus at 0°, they were not located at a trough or a peak. Contrast this failure with test 43, in which the ant also reached zero in steps, but the zero plateau is in a trough. The ant faced both goals in tests 26, 33 and 39. Test 43 is a rare example of the ant actually reaching the goal. Ant L faced the real goal in tests 38 and 47, the fictive goal in tests 26 and 33, both goals in tests 14 and 33. There were three failures (tests 22, 38, 39).

Table 1 summarises the proportion of successes across the different tests of the nine tests. The early tests given before any training were as successful as those given after training thus indicating that the memory of a route direction persists overnight. The success rate does not change appreciably over the course of the experiment. If we take each test as an individual data point and assume that the probability of a success is 40/360 (i.e. facing ±10° at the real or fictitious goal, then the frequency of successes is much greater than would be expect by chance (p<0.000001 binomial test, for 80/360 binomial test is p=000447).

**Table 1:** Number of successes in the tests administered to 9 ants

| Ant | No. tests- no. successes | Test: | No. ants-successes |
| --- | --- | --- | --- |
| A | 8-3 | 14 | 7-5 |
| B | 7-4 | 22 | 7-3 |
| E | 9-6 | 26-early | 9-6 |
| L | 9-7 | 30 | 9-4 |
| M | 6-4 | 33-early | 9-7 |
| O | 8-5 | 38 | 8-5 |
| Q | 9-4 | 39-early | 7-2 |
| T | 6-4 | 43-early | 7-3 |
| U | 7-3 | 47 | 6-5 |
| **Totals** | 69-40;<br>Success/total=0.58<br>31 trials to real goal<br>19 trials to fictive goal | Early tests: 32-18;<br>Success/ total= 0.56<br>Other tests: 37-22;<br>Success/total=0.59 |  |

Most paths do not reach either the real or the fictive goal. Nonetheless, some ants tend to travel for a short segment in a relatively consistent direction from the start point. To determine the distribution of directions over a segment of the path beginning at the start point, we selected suitable endpoints by eye for each of the 69 tests. The duration of the segments in Figure 4 emphasises how short some of the samples are. We then calculated the distance travelled in different orientations as described in the Methods. Since ants often move in two opposite directions, the plots show the orientation of the axis of travel relative to the expected magnetic axis, rather than travel direction. The distribution of distances travelled along each segment usually has one or two clear peak orientations, as shown by the distributions for each recorded test from ant L (Figure 4). The peak orientations in some tests are close to the magnetically defined axis (0°). Others are oblique to or across the axis.

**Figure 4.**
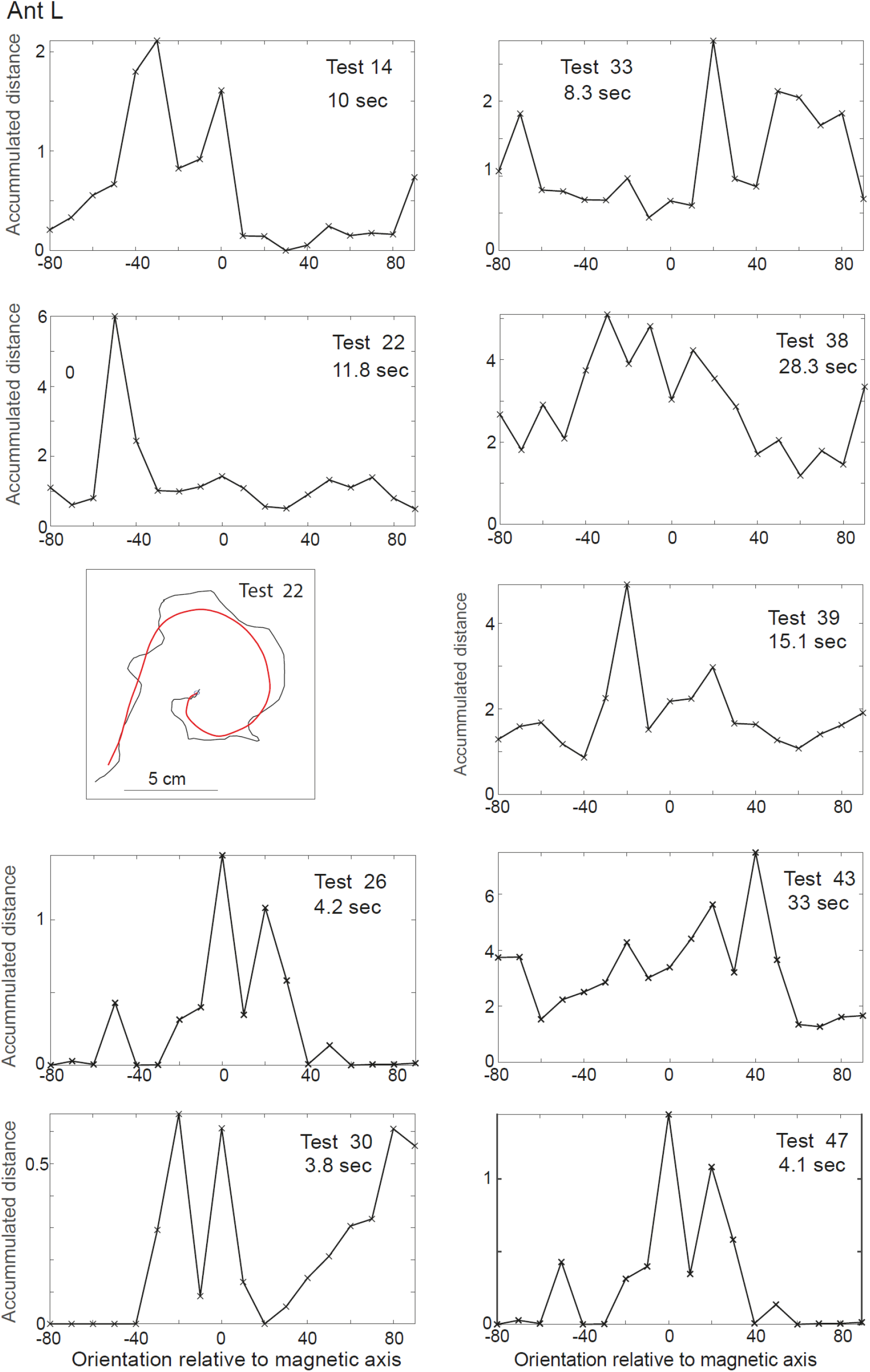
Path orientations of ant L. Each panel shows the distribution of path orientations along a selected segment of one test. X axis is the axial orientation relative to the specified magnetic axis. Y axis is the accumulated distance within each 10° bin of directions. The panel below the test 22 histogram shows the unsmoothed (black) and smoothed (red) paths

We accumulated the peak value of each individual test distribution, as in Figure 4, across all 69 tests from 9 ants. The resulting distribution (Figure 5) is quite broad, but with a peak close to zero, suggesting a weak tendency for the ants path to be oriented along the magnetically defined axis, with additional components across the axis.

**Figure 5.**
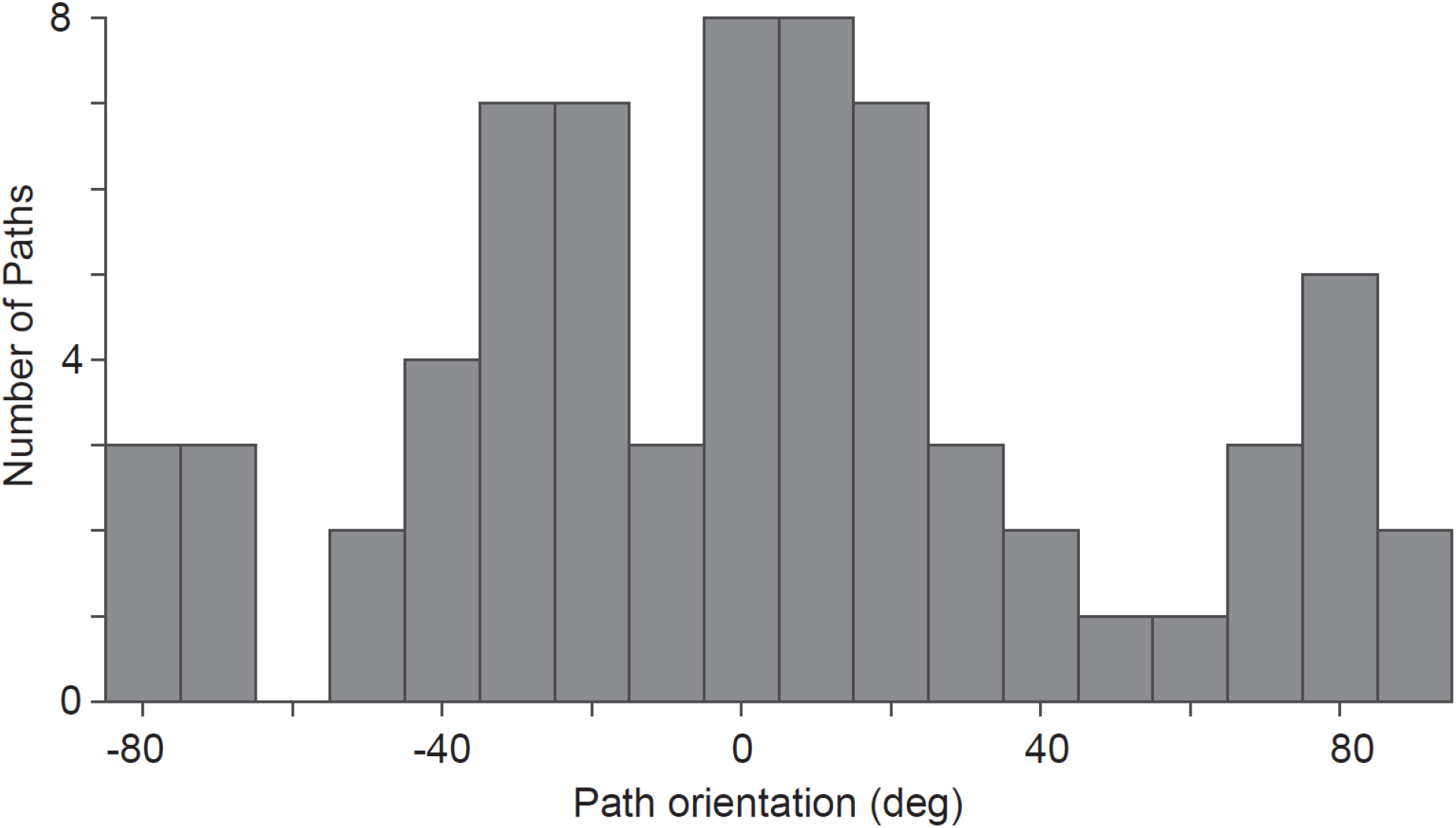
Distribution of the peak orientations from all the tests of 9 ants. Bin width 10°. Mean of distribution: 2.9°, standard deviation: 43.8°, n: 69.

#### (iii) Room cues to direction

The results so far emphasise that, in addition to moving towards the goal, ants often face and move in the opposite direction during training and tests trials. One of several possible reasons for this unexpected behaviour is the presence of competing directional cues that are fixed to the room, rather than to the rotating magnetic direction. For instance, it is unlikely that the illumination of the arena is symmetrical. To examine the possibility that there are competing directional cues, we ran a special test at the end of the experiment. We gave the ants three consecutive training trials in which the magnetic field was in the same direction relative to the room and to any cues associated with it. This training was followed by a test in which magnetic direction was rotated 90° clockwise from the training direction. The ants’ idiosyncratic paths in the test were in both the previous training direction (marked ‘room’) and in the magnetically specified test direction, at about 11 o’clock (Figure 6). Moments in the path when the ant faced in the room specified direction (±10°) are marked by red disks.

**Figure 6.**
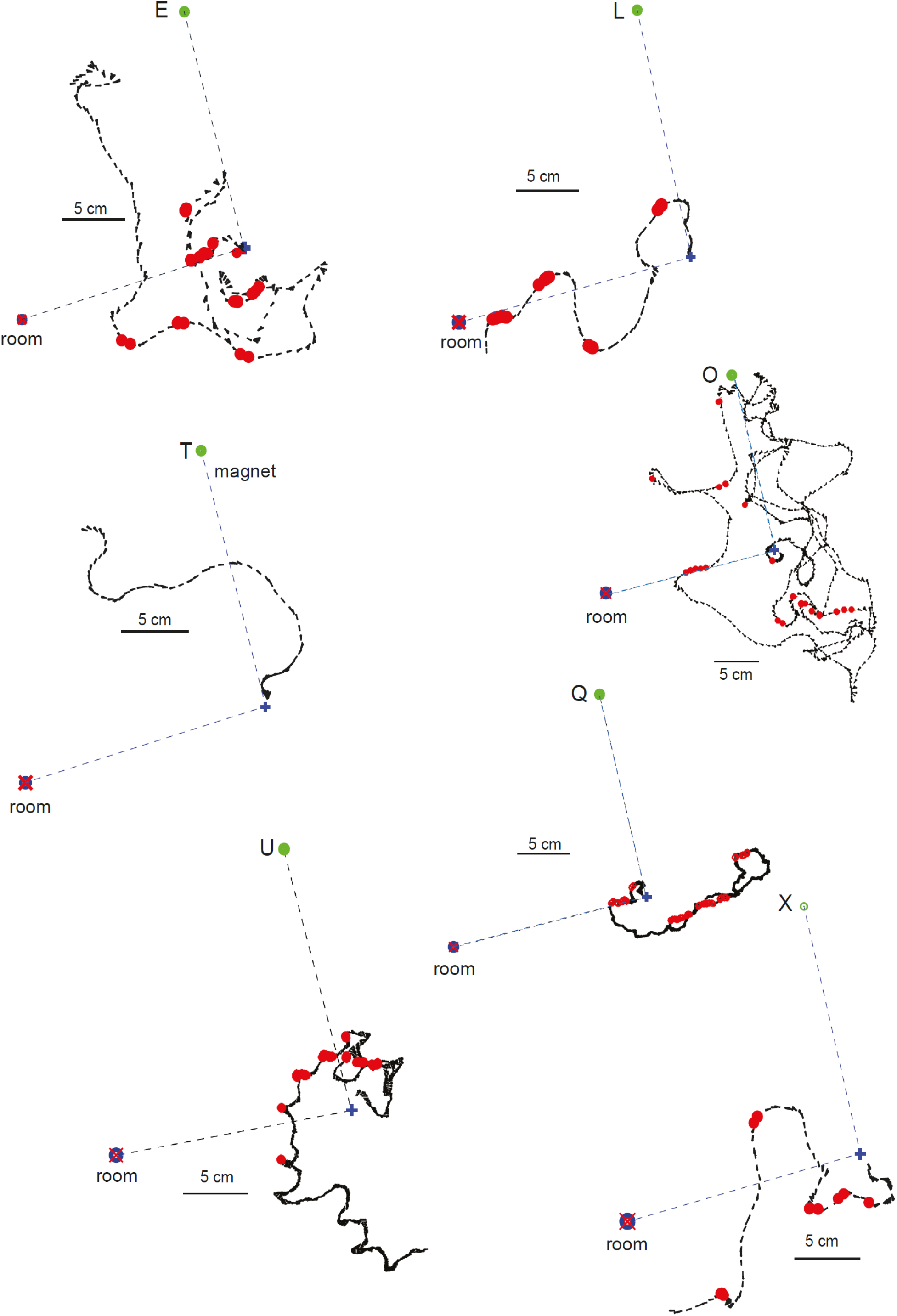
Test after three consecutive training trials with food in the same direction relative to the room. Training direction is indicated by a marker at the goal distance labelled ‘room’. Red disks show when ant faces the marker within ±10°. The paths of all tested ants are included. Other details as in Figure 1.

The ants followed a variety of paths (Figure 6). Ants E and Q moved in the room direction at or near the start of the path. Ant Q’s subsequent path was in the diametrically opposite direction followed by a long section towards the room specified goal. Ant L differed; it began with a stretch in the magnetically defined direction and then nearly reached the room specified goal position. Ant O took a roundabout route to the magnetically defined goal and, then on its way to the fictive goal, it had a clear stretch towards the room defined goal. Ant X began in the opposite direction from the magnetic goal and then moved towards the room goal before wandering off. Ant T began with a short stretch towards the magnetic goal, and ignored the room specified goal. Ant U’s intentions were unclear. It took a semi-circular path around the nest during which briefly changed direction and faced towards the room goal. Thus, 4 (E, L Q, X) out of the 7 tested ants had clear path segments towards the room goal. The precision of these paths is too great to be attributable to the kind of broadly distributed cross-axis paths seen in Figures 4 and 5. Though insufficient as proof, this single test suggests that the ants may be unwilling to rely entirely on magnetic cues and are eager to find other more reliable directional cues (Johnsen et al., 2020).

### 2. Can ants use magnetic direction as a contextual cue?

In this experiment, the ants were again released at the centre of the arena. They were confronted with an upright and an inverted triangle in fixed positions, 90° apart, on the cylindrical arena wall with their bases close to the floor (Figure 7A). When the magnetic direction from the central start point to the upright triangle was North, sucrose was placed below the upright triangle. Sucrose was below the inverted triangle when the magnetic direction to that triangle was West (Figure 7A). Thus, ants can only learn the task if they can appreciate that there is a linkage between the orientation of the triangle and the direction of the magnetic field from the centre to the triangle. Formulated In ecological terms, the problem is to go in one of two directions to one of two feeding sites with the choice set by what can be seen there.

**Figure 7.** Training to two triangles. **A.** Schematic of training arrangement showing magnetic directions from centre of arena to base or apex of triangle. Cross shows which triangle is rewarded. **B.** Left: Proportion of trials (training and tests) in which the upright triangle was reached first. Dashed line trials with upright triangle rewarded; solid line inverted triangle rewarded. Right performance of individual ants. Proportion of trials in which upright triangle was reached vs. proportion of correct trials. **C**. Individual trajectories on trial 60 – a training trial. Position of triangles shown in yellow. Trajectory coloured red when ants face one or other triangle within ±10°.

Despite many training trials, ants did not learn to approach the correct triangle. During the course of training and testing, individual ants were recorded on ten trials between trials 27 and 60. For each recorded trial, we noted which triangle the ants reached first, ignoring the two times in which an ant failed to reach either triangle. We then plotted the proportion of ants approaching the upright triangle (Figure 7B left). Irrespective of which triangle was signalled, ants tended to go to the upright triangle with no improvement during the course of training.

The performance of individual ants was assessed using 14 individuals that were recorded on 5 or more of the recorded ten trials. For each ant, we plotted the proportion of trials in which the ant first reached the upright triangle against the proportion of trials in which the ant first reached the correct triangle. 9 out of 14 ants chose incorrectly on half or more of their recorded trials. 11 out of 14 ants preferred to approach the upright triangle (Figure 7B right).

The uncertainty of the ants’ behaviour is illustrated through individual paths taken from the last training trial (60) in which the upright triangle was rewarded (Figure 7C). Whichever triangle the ants reached first, they tended to approach the non-chosen triangle as well. Ants sometimes just switched their path from one triangle to the other (Ants B, Z, I). At the extreme, they oscillate between the two over several cycles (Ant F). Despite this indecision, the ants were still strongly biased in favour of reaching the upright triangle first. This bias could be because the upright triangle is more ecologically plausible and that the visual system is better tuned to it.

In another experiment of the same kind, the triangles were placed 180° apart. In this case there was a weaker preference for the upright triangle (inverted triangle 17/59 correct choices, upright triangle: 22/41 correct choices, Two tail Fisher exact probability test: P= 0.014). Again there was no evidence that ants learnt to choose the correct triangle.

## Discussion

The major contribution of this study is to provide evidence that ants can learn and remember overnight the magnetic direction of a route between a starting point and a feeding place. Behavioural signs of this memory tend to be restricted to the very start of the route. At least two factors could contribute to this short distance. One is the small area over which magnetic direction could be reliably manipulated. At the start of normal routes in larger spaces, the route vector and desired compass direction coincide for a while. In a small arena, slight deviations from the route will cause the direction of the route vector to diverge from the specified magnetic direction, so confusing the ant over the correct path to follow. A second reason for leaving the magnetically defined route is the existence of competing cues from other directional signals (Figure 7). The ants may be unsure whether they should move in the specified magnetic direction or follow another cue.

### Is moving in the opposite direction to a goal a sign of uncertainty?

A second feature of the ant’s routes is the frequency with which ants face or move in precisely the opposite direction from the magnetically signalled route. Both directions often occur in the same path (Figures 2 and 3).

In a sense, it should be no surprise that ants switch between opposite directions. Inverting route vectors between their nest and a food site often occurs in bees and ants. And a switch in roue direction can be induced by just starving or feeding bees (Dyer et al., 2002) or ants (Harris et al., 2005).

Taking the opposite path to the expected direction or ‘back-tracking’ (Schwarz et al. 2011, Wystrach et al. 2013) is known to occur in ants which are faced with an unexpected situation or are thwarted from reaching a goal. The various benefits of back-tracking have been articulated elsewhere as returning to familiar terrain when the scene is unfamiliar, so providing an opportunity for correcting errors in an approach (Collett et al., 1993, Wystrach et al 2013), or for choosing a new approach to a goal (Wystrach et al. 2020, M. Collett, unpublished data), or for learning a route in the reverse direction (Graham and Collett, 2006). Back-tracking may also underlie the evolution of the learning walks and learning flights that ants bees and wasps perform when they first leave their nest (Collett and Zeil, 2018; Zeil and Fleischman, 2019). During these manoeuvres the insects look back multiple times to face their nest and learn what the immediate surroundings of the nest look like. Looking back might originally be a response to experiencing views of unfamiliar scenery that an insect encounters on first emerging and moving a short distance away from its nest.

These circumstances do not apply in the present case. In both route learning and route selection, the ants’ behaviour seems rather a response to their uncertainty about the correct travel direction. This possibility is emphasised by the second experiment in which ants were uncertain about which triangle to choose. Should they approach the magnetically indicated or the more attractive triangle? The undecided ant may not know the location of the triangle that it is not currently approaching and by turning away, it unlocks its attention and frees it to gain a new view of the scene. It can then switch its path towards the other triangle. This kind of behaviour was first described decades ago in walking, wingless *Drosophila*. The flies, when confined to a small arena with two inaccessible visual targets, oscillate between them for several hours, like Buridan’s fictional ass trying vainly to choose between two similar piles of hay (Bülthoff et al, 1982).

## Acknowledgements

TSC thanks Peter Reed for electronic support and the Leverhulme Trust for funding through an Emeritus Fellowship (EM-2016-066). AP was funded by EPSRC grants EP/P006094/1 and EP/S030964/1.

## Supplement

**Supplementary Figure 1.**
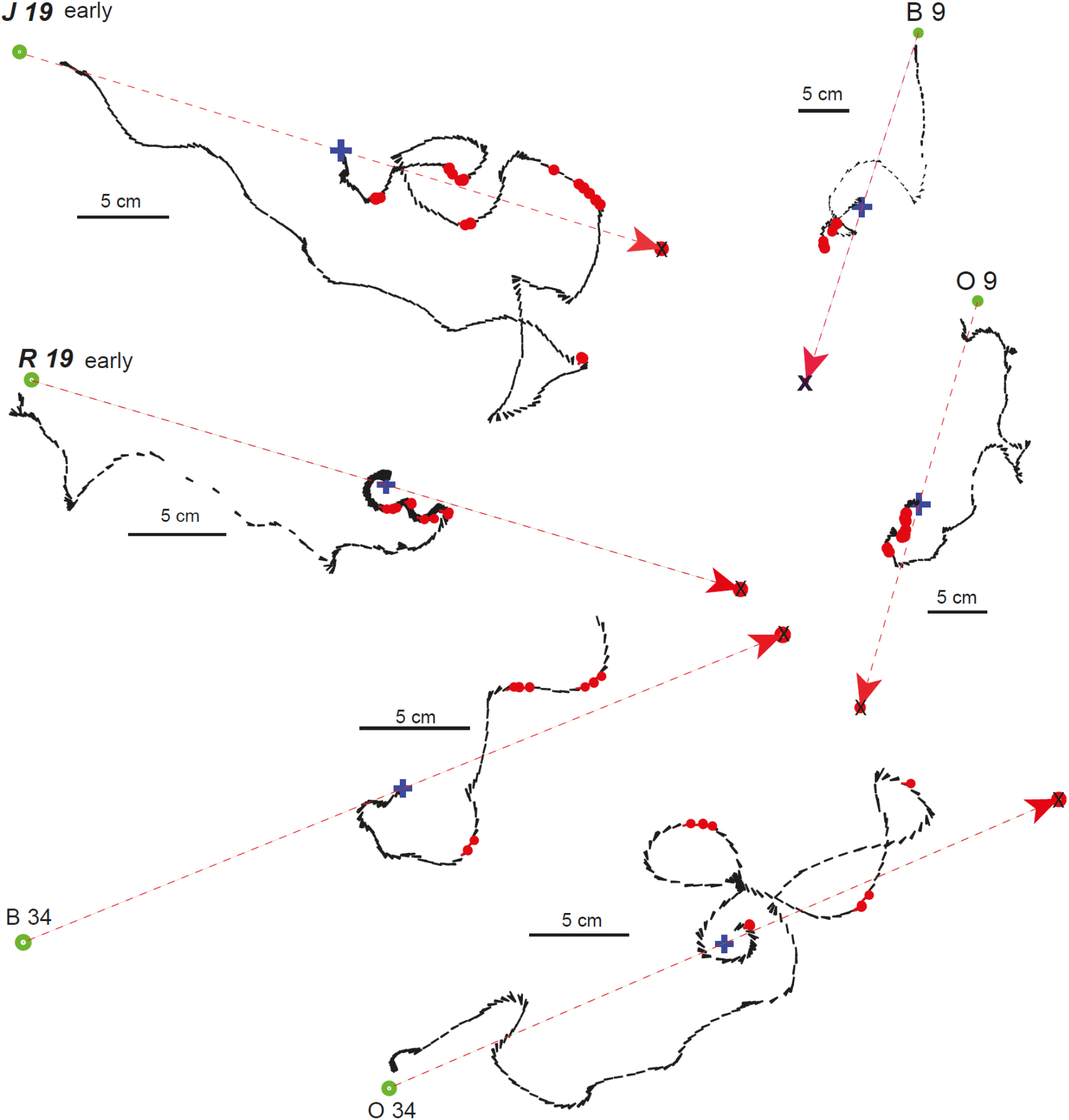
More training trials with magnetic direction supported by a black bar. Each panel shows the path of one ant on different training trials with the position and orientation of the ant shown at 30 fps. Blue cross marks starting point. Red disks show when the ant faces within ±10° either towards the food or a virtual goal 180° in the opposite direction. The marker with an X shows the position of the virtual goal and the other marker the food. The arrow emanating from the start point sets whether the red dots indicate the ant facing the real or virtual goal. Arrows are blue when the red dots mark facing towards the real goal. Arrows are red when the red dots indicate facing towards the virtual goal. In trials marked early, the coil system was off and the ant was guided by the Earth’ magnetic field.

